# More accurate estimation of cell composition in bulk expression through robust integration of single-cell information

**DOI:** 10.1101/2022.05.13.491858

**Authors:** Ali Karimnezhad

## Abstract

The rapid single-cell transcriptomic technology developments has led to an increasing interest in cellular heterogeneity within cell populations. Although cell-type proportions can be obtained directly from single-cell RNA sequencing (scRNA-seq), it is costly and not feasible in every study. Alternatively, with fewer experimental complications, cell-type compositions are characterized from bulk RNA-seq data. Many computational tools have been developed and reported in the literature. However, they fail to appropriately incorporate the covariance structures in both scRNA-seq and bulk RNA-seq datasets in use.

We present a covariance-based single-cell decomposition (CSCD) method that estimates cell-type proportions in bulk data through building a reference expression profile based on a single-cell data, and learning gene-specific bulk expression transformations using a constrained linear inverse model. The approach is similar to Bisque, a cell-type decomposition method that was recently developed. Bisque is limited to a univariate model, thus unable to incorporate gene-gene correlations into the analysis. We introduce a more advanced model that successfully incorporates the covariance structures in both scRNA-seq and bulk RNA-seq datasets into the analysis, and fixes the collinearity issue by utilizing a linear shrinkage estimation of the corresponding covariance matrices. We applied CSCD to several publicly available datasets and measured the performance of CSCD, Bisque and six other common methods in the literature. Our results indicate that CSCD is more accurate and comprehensive than most of the existing methods.

## 1 Introduction

Gene expression profiling measures mRNA levels, showing the pattern of genes expressed by a cell at the transcription level (Fielden and Zacharewski 2001). It is widely practised by a variety of biomedical researchers, molecular biologists and environmental toxicologists, to characterize cellular or disease states (Tsoucas *et al*. 2019).

RNA-sequencing of the individual cell or single-cell RNA-sequencing (scRNA-seq) has enabled dissecting transcriptomic heterogeneity, and led to the discovery of novel cell types (Zheng *et al*. 2017). The rapid single-cell transcriptomic technology developments has led to an increasing interest in cellular heterogeneity within cell populations. Cell-type compositions may be characterized from bulk RNA-seq data, which typically represent total gene expression levels of heterogeneous cell types within tissues.

Many computational tools have been developed in the literature for dissecting cellular content from bulk samples. Perhaps the non-negative least-squares (NNLS) and ordinary least squares (OLS) methods are the most simple approaches utilized in the cell-type decomposition literature (Abbas *et al*. 2009, Avila *et al*. 2020). However, as we show in our data analysis, they may either fail or lead to poor performance. CIBERSORT (Newman *et al*. 2015) and its next generation, CIBERSORTx (Newman *et al*. 2019), extract cell-type specific gene expression signatures from a reference scRNA-seq dataset. They learn cell-type decomposition in bulk expression data by applying a linear support vector regression (SVR) method. These machine learning-based tools were originally designed to analyze microarray data and are expected to perform well for purified cell populations. BSEQ-sc (Baron *et al*. 2016) extends CIBERSORT and allows for building a reference profile based on scRNA-seq gene expression. Similar to the above methods, the dampened weighted least squares (DWLS) approach of Tsoucas *et al*. (2019) uses scRNA-seq data to extract cell-type specific gene expression signatures, and estimates cell-type proportions by improving the OLS approach. The corresponding tool also outputs cell-type proportions estimated by the OLS and SVR approaches based on the generated signature matrix. Note that because the signature matrices created by DWLS and CIBERSORTx are different, the SVR approach used in both tools leads to distinct estimates for the same cell-type proportions.

MuSiC (Wang *et al*. 2019) performs cell-type decomposition without the need to build a gene expression signature matrix. It utilizes gene expression levels in the scRNA-seq reference data and applies a constrained NNLS regression model to estimate cell-type proportions in the bulk expression data. The corresponding tool also outputs estimated cell-type proportions using the ordinary NNLS method. MuSiC was shown to outperform NNLS, CIBERSORT and BSEQ-sc on two real datasets.

The above-mentioned approaches do not apply any transformation to either scRNA-seq or bulk RNA expression datasets, and assume that there is a direct proportional relationship between both datasets. Bisque (Jew *et al*. 2020) highlights that different technologies used to generate bulk and scRNA-seq data may cause an issue for decomposition models and the direct proportional relationship assumption may not be true. It was shown in the corresponding publication that these differences may introduce gene-specific biases that negatively impact the correlation between cell-type-specific and bulk tissue measurements. It was concluded that expression profiles of cell types in heterogeneous tissues may not follow the direct proportionality assumptions of regression-based methods. By accounting for such biases, Bisque estimates cell-type proportions in bulk expression data through robust integration of single-cell information, and uses a normalizing transformation on both bulk and single-cell expressions. Bisque was applied on two datasets in which both bulk and single-cell expressions were collected from the same individuals. It was shown that Bisque outperforms CIBERSORT, CIBERSORTx, MuSiC and BSEQ-sc. However, Bisque does not account for gene-gene correlations. The normalization in Bisque is done for each gene separately, while multivariate normalization that accounts for gene-gene correlations seems to be more appropriate. Indeed, to the best of our knowledge, none of the existing methods in the literature have incorporated a multivariate normalization technique to provide such such useful information into the analysis, perhaps due to the existence of high gene-gene correlations which in turn leads to the collinearity issue. Here, we introduce CSCD, an advanced model for analyzing bulk RNAseq datasets that improves on the Bisque method. It estimates cell-type proportions more accurately and successfully utilizes gene-gene correlations into the analysis. We fix the collinearity issue by utilizing a linear shrinkage estimation of the corresponding covariance matrix. We apply CSCD to several publicly available datasets in which both bulk and scRNA-seq were collected from the same individuals. Using gene expression levels in both bulk RNAseq and scRNA-seq data for the same individuals maximizes measurement accuracy, as the real cell-type proportions are known in advance. We compare the performance of CSCD with Bisque, CIBERSORTx, MuSiC, DWLS, OLS, NNLS and SVR. Although our primary aim is to improve on Bisque, our results indicate that CSCD is more accurate and comprehensive than most of the other existing methods used in this paper, as well.

The structure of this paper is as follows. In Section 2, we introduce the notations used in this paper, and explain the method in detail. Our proposed method is built on the model used in Bisque and we quickly review the workflow of Bisque. In Section 3, we evaluate the performance of CSCD, Bisque, CIBERSORTx, MuSiC, DWLS, OLS, NNLS and SVR using several publicly available datasets. Discussion and concluding remarks are provided in Section 4.

## 2 Methods

### 2.1 Notation

Let *C, J* and *I* represent cell types, subjects in the single-cell data and subjects in the bulk expression data, respectively. Also let *G* represent the total number of genes common in both bulk and single-cell data. For each gene *g* ∈ {1, …, *G*} and cell type *c* ∈ {1, …, *C*}, we denote average relative abundances across all subjects in the single-cell dataset by *Z*_*gc*_. For each cell type *c* and subject *j* ∈ {1, …, *J* }, let *p*_*cj*_ represent the fraction of cell type *c*. For each gene *g* and subject *i* ∈ {1, …, *I*}, we denote counts per million (CPM) converted relative abundances in the bulk data by *X*_*gi*_.

Given *p*_*cj*_ and *Z*_*gc*_, the pseudo-bulk for each gene *g* and subject *j* is given by 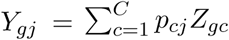, which is a weighted sum of average relative abundances. Let ***Y*** = [*Y*_*gj*_] be the matrix containing all *Y*_*gj*_ with genes in rows and subjects in columns. Also, let ***X*** = [*X*_*gi*_] represent the bulk expression data matrix. The corresponding sample mean vectors are given by 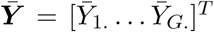 and 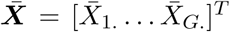, with 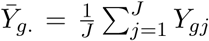 and 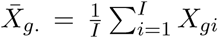. The sample covariance matrices associated with the bulk and single-cell expression datasets are then respectively given by

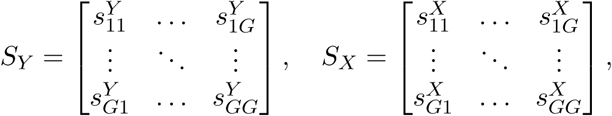

where for 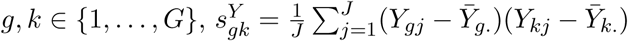 and 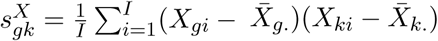.

### 2.2 A novel method

Bisque estimates cell-type proportions *p*_*ci*_ in bulk RNA-seq data through the integration of single-cell information. In a pre-processing step, Bisque excludes genes with zero variance in the single-cell dataset, unexpressed genes in the bulk expression and mitochondrial genes. Since the distribution of bulk expression (*X*_*gi*_) may differ from the distribution of the single-cell expression (*Y*_*gj*_), Bisque re-scales the bulk expression data so that the two distributions match more closely. This maximizes the linear relationship across all genes for improved decomposition. Bisque replaces *X*_*gi*_ by the gene-specific transformation 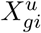 so that it satisfies 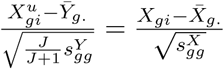, where 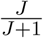 is a variance shrinkage factor, due to the small number of subjects in the single-cell dataset. Then, for each subject *i* in the bulk expression data, cell-type proportion *p*_*ci*_ is estimated using the NNLS regression with a sum-to-one constraint, i.e., they minimize the norm

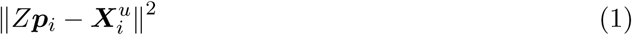

where

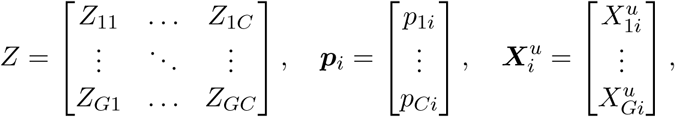

such that 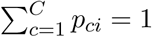.

We apply the same pre-processing steps used in Bisque. Although the univariate transformation *X*^*u*^ in Bisque accounts for changes across individuals in both the bulk and single-cell datasets, it is unable to account for gene-gene correlations. Consequently, this may affect the estimation accuracy. Instead of the transformation 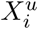, we suggest replacing ***X*** by ***X*** ^*m*^ so that it satisfies 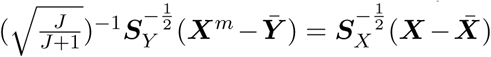, or equivalently 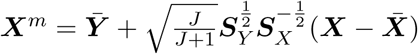, where 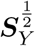 and 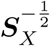 stand for the root of the sample covariance matrix ***S***_*Y*_ and the inverse of the root of the sample covariance matrix ***S***_*X*_, respectively.The above transformation leads to inclusion of gene-gene correlations into the model. However, calculating the inverse of and/or the root of the covariance matrices appeared in the transformation may be a challenge. Gene expression covariance matrices have a very large dimension, compared to the number of observations/subjects. It is well known that such covariance matrices perform poorly, are expected to suffer fromill-conditioning, and might not even be invertible. To overcome such challenges, there are ways to arrive at modified covariance matrix estimators. Incorporating some additional knowledge in the estimation process, such as sparseness, a graph model ora factor model may be ideal, see for example Bickel and Levina (2008), Khare and Rajaratnam (2011) and Cai and Zhou (2012) among many others.However, such additional knowledge is not always available. Alternatively, it may be reasonable to look for rotation-equivariant covariance matrixestimators, i.e., matrices with the same eigenvectors. This implies that a sample covariance matrix can be differentiated by its eigenvalues.On the other hand, according to Ledoit and Wolf (2004), the largest sample eigenvalues are biased upwards, and the smallest ones downwards. Therefore, it is possible tomodify the sample covariance matrix by shrinking its eigenvalues towards their grand mean. Indeed, shrinkage estimation of the covariance matrix has become one of the most successful approaches. Ledoit and Wolf (2004) introduce a linear shrinkage estimation of the covariance matrix, and later, Ledoit and Wolf (2012) present a non-linear shrinkage estimation method.The latter clarifies that if the number of variables (genes in the current study) are significantly higher than the number of observations, the linear shrinkage estimation is more appropriate than the non-linear approach. To fix the above-mentioned issue, we apply the linear shrinkage estimation of Ledoit and Wolf (2004) on the sample covariance matrices ***S***_*Y*_ and ***S***_*X*_, and denote the resulting estimates by ***S***_*Y,s*_ and ***S***_*X,s*_, respectively. The final transformed bulk expression data matrix is then given by

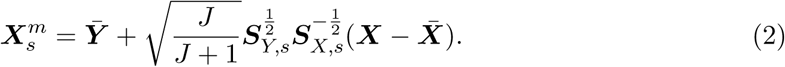

Cell-type proportions can now be obtained by minimizing the norm (1), with 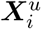 being replaced by the *i*-th column of 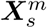. The covariance-based bulk expression transformation 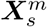 in (2) is ideal due to the fact that it is able to extract and utilize all the information (univariate and multivariate changes) from both bulk and single-cell datasets into analysis. Nevertheless, the inclusion of all genes to the analysis may lead to imprecise results. Obviously, if one categorizes all genes to several cell-type clusters, some genes may appear in all or most of them. This may negatively influence the cell-type proportion estimation accuracy. To overcome this issue, we identify cluster marker genes (marker genes that are recognized to be differentially expressed within the cell-type clusters) using the `FindAllMarkers` function of the `Seurat` package (Hao *et al*. 2021). In this approach, the expression for a given gene *g* among cells in a given cell-type cluster *c* is compared against the expression of that gene among cells in all other cell-type clusters, say 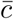. We look at the percentage of cells in the cluster *c* where the gene is detected (say *p*^*gc*^) and the percentage of cells of other types (say 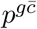). Ideally, a cluster marker gene is expected to be expressed exclusively in that cluster and silenced in all other clusters and thus for each gene *g* in cluster *c, p*^*gc*^ is expected to be high (towards one) and 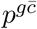 is expected to be low (towards zero). It may be reasonable to consider a 0.5 threshold for the difference between *p*^*gc*^ and 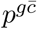. Thus, we excluded marker genes for which 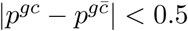. The 0.5 threshold may also be replaced by some arbitrary values. The smaller this threshold, the more genes to be included in the final list of gene markers. We further remove common marker genes that appear in multiple clusters. This leads to a list of final marker genes that each falls into only one cell-type cluster and reduces the number of common genes between cell types to zero. The final gene list is then used to compute the transformed expression levels 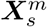 in (2). The final cell-type proportions ***p***_*i*_ are given by minimizing the norm (1) with 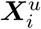 being replaced by the *i*-th column of 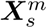.

## 3 Results

The reference-based decomposition approach presented in this paper requires both bulk and scRNA-seq datasets that contain expression levels of multiple subjects. ScRNA-seq data may be collected from individuals involved in the same study or similar studies conducted by other labs. It is assumed that scRNA-seq data includes cell-type as well as subject labels. A reference profile (*Z*) is constructed by read count abundance averaging within each cell-type category. Given the generated single-cell profile and cell-type proportions observed in the single-cell data, the method applies a multivariate transformation to the original bulk data. Given the single-cell reference profile and transformed bulk expression data, CSCD learns cell-type composition using a constrained linear inverse model (Wang *et al*. 2017).

### 3.1 Performance Evaluation

In this section, we evaluate the performance of CSCD and seven other common tools (Bisque, CIBERSORTx, MuSiC, DWLS, OLS, NNLS and SVR) on several publicly available datasets. Table 1 represents distribution of different cells in the single-cell datasets used in this paper. Accuracy was maximized by using datasets for which both bulk and single-cell expression levels were available from the same individuals. Thus, real cell-type portions for all samples in the bulk expression data are known. The Spearman correlation (R) has become a common tool in measuring cell-type estimation accuracy in the literature. In addition to reporting R, we measure the difference between the real and estimated cell-type proportions by mean squared error (MSE) and mean absolute error (MAE) to evaluate the performance of different methods. The higher the correlation R, the better the model performance. Similarly, the smaller the MSE (or MAE), the more accurate the corresponding method.

**Table 1:** Distribution of cells in different single-cell datasets used in this paper.

| Dataset | Cells and counts |  |  |  |  |  |  |  |  |  |  |  |  |
| --- | --- | --- | --- | --- | --- | --- | --- | --- | --- | --- | --- | --- | --- |
|  | neuronal progenitor cell (NPC) | definitive endoderm cell (DEC) |  |  | endothelial cell (EC) | trophoblast-like (TB) |  |  | human foreskin fibroblast (HFF) |  |  | H1 | H9 |
| human embryonic stem cells | 173 | 138 |  |  | 105 | 69 |  |  | 159 |  |  | 212 | 162 |
| pancreatic islets data | pluripotent stem cell (PSC) | acinar | alpha | beta | co-expression | delta | dacutal | endothelial | epsilon | gamma | unclassified endocrine | unclassified cell |  |
|  | 18 | 86 | 254 | 101 | 17 | 21 | 89 | 9 | 4 | 48 | 9 | 1 |  |
| cortex tissue data | astrocyte (Astr) | endothelial (Endo) | excitatory neuron (Exc) | inhibitory neuron (Inh) | microglia (Micr) | oligodendrocyte progenitor cell (OPC) |  |  | oligodendrocyte (Olig) |  |  | pericyte (Peri) |  |
|  | 10256 | 876 | 19150 | 8233 | 2496 | 2586 |  |  | 16232 |  |  | 559 |  |

### 3.2 Evaluation on human embryonic stem cells

We first benchmarked all the eight decomposition methods using expression data collected from 1018 purified human embryonic stem cell samples of seven cell types generated from various lineage-specific progenitors (Chu *et al*. 2016). See Table 1 for the distribution of the cells in the corresponding single-cell reference data. A total of 19 bulk samples from these cell types were sequenced. This includes duplicates for DEC, NPC, and TB, triplicates for H9, EC, and HFF, and four replicates for H1. Both bulk and scRNA-seq data were collected from the same individuals. For more accuracy, we merged replicates of different cell types which reduced the bulk data to seven samples. Here, with a different purpose than Chu *et al*. (2016), we analyzed the bulk and single-cell expression data to assess the accuracy of the different cell-type decomposition methods.

Figure 1 represents a jitter plot of the estimated cell-type proportions using the different methods for the seven purified bulk samples. Each panel represents real and estimated values for the labelled cell type. Because cell samples are purified, only one individual in each panel is expected to have a cell-type proportion of one and the remaining cell-type proportions should be zero. The figure represents proportions estimated by CSCD, Bisque, CIBERSORTx, SVR, OLS and DWLS. MuSiC and NNLS failed due to a “not enough valid cell type” error. Based on the real and estimated cell-type proportions, it is obvious from the jitter plot that CIBERSORTx and SVR performed very well. This was expected, as CIBERSORTx was originally developed for data that utilizes a reference generated from purified cell populations, see Jew *et al*. (2020). The SVR approach implemented in DWLS also uses a similar approach to estimate cell-type proportions. Based on the figure, DWLS seems to be the third top method. The performance of these tools is compared in Table 2, which summarizes the R, MSE and MAE measurements. The results in the table also confirm CIBERSORTx and SVR methods significantly outperformed the other methods. DWLS also outperformed the rest of the methods. Although Bisque led to a bit higher correlation than CSCD, it led to higher MSE and MAE values than CSCD. In two of the three measurements, CSCD outperformed Bisque.

**Table 2:** Performance measures of different cell-type estimation methods based on seven bulk samples in the human embryonic stem cells data. MuSiC and NNLS failed to estimate cell-type proportions.

| Measure | CSCD | Bisque | CIBERSORTx | MuSiC | SVR | NNLS | DWLS | OLS |
| --- | --- | --- | --- | --- | --- | --- | --- | --- |
| R | 0.6124 | 0.6513 | 0.7203 | — | 0.7445 | — | 0.6042 | 0.5823 |
| MSE | 0.0245 | 0.0321 | 0.0047 | — | 0.0058 | — | 0.0123 | 0.1109 |
| MAE | 0.1051 | 0.1063 | 0.0281 | — | 0.0179 | — | 0.0361 | 0.1813 |

**Figure 1:**
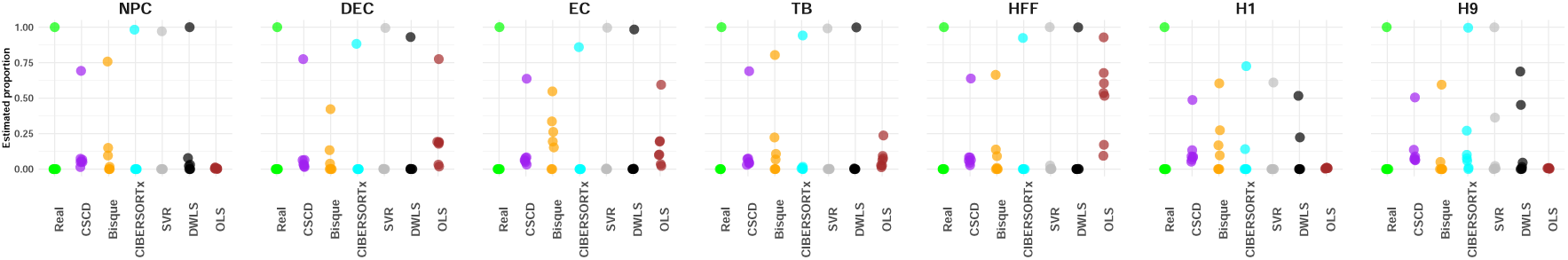
Jitter plot of estimated cell-type proportions for seven bulk samples in human embryonic stem cells data. Each panel represents estimated values for the labelled cell type. Different colors represent different estimation methods. MuSiC and NNLS failed to estimate cell-type proportions.

### 3.3 Evaluation on pancreatic islets data

We also benchmarked these decomposition methods using expression data collected from human pancreatic islets (Segerstolpe *et al*. 2016). The scRNA-seq data were collected from six healthy and four type-2 diabetic (T2D) donors. The bulk data were collected from three of the six healthy individuals in the scRNA-seq data and four different T2D donors.

To maximize measurement precision, we evaluated the performance of the different methods based on the three healthy donors for which both bulk and scRNA-seq data are available. See Table 1 for the distribution of the cells. We excluded the unclassified cell from our analysis.

Figure 2 represents a jitter plot of the estimated cell-type proportions using the different methods for the three common donors. Different methods performed differently. It is not easy to make an overall judgment based on this figure alone. However, the figure is informative in the beta estimates, which was reported to be very important (Wang *et al*. 2019). The real beta proportions in the scRNA-seq data are around 20 percent. Also, CSCD was the only method that correctly detected real beta proportions in the bulk samples. For a more quantitative comparison, Table 3 summarizes the R, MSE and MAE values. The table illustrates that all methods led to positive correlation, with CSCD gaining the highest correlation. Looking at the MSE and MAE measurements, we observe that CSCD outperformed all the other methods. Bisque and SVR were the second and third top methods leading to minimum MSEs and MAEs.

**Table 3:**
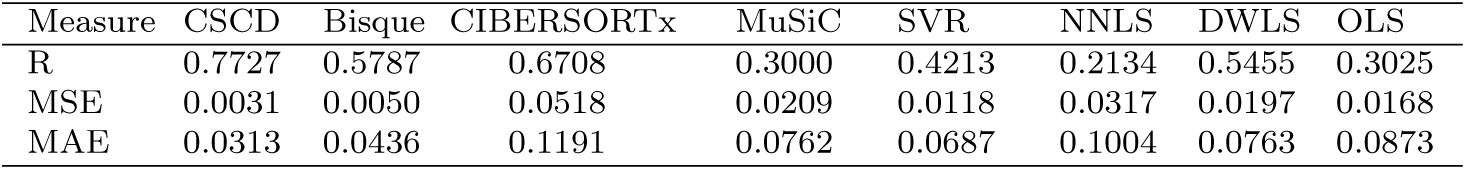
Performance measures of different cell-type estimation methods based on three bulk samples in the pancreatic islets data.

**Figure 2:**
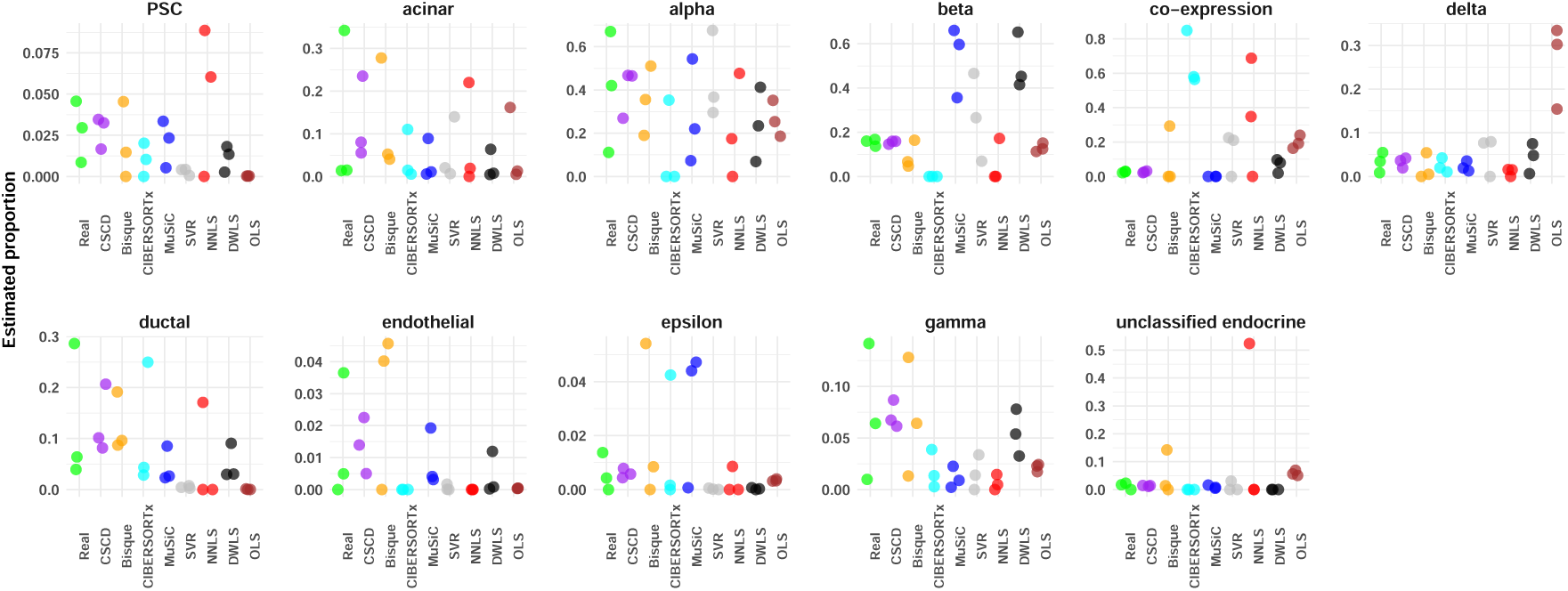
Jitter plot of estimated cell-type proportions for three bulk samples in the pancreatic islets data. Each panel represents estimated values for the labelled cell type. Different colors represent different estimation methods.

Cell-specific decomposition accuracy was also evaluated using two other well-characterized datasets. Fadista *et al*. (2014) provide a comprehensive catalog of novel genetic variants influencing gene expression and metabolic phenotypes in human pancreatic islets. Their study includes expression levels of 89 bulk donors. The data also includes hemoglobin A1c (HbA1c) levels, which is an important biomarker for T2D and is expected to have a negative relationship with beta proportions (Wang *et al*. 2019). Indeed, T2D is characterized by the inability of the beta cells to secrete enough insulin to overcome insulin resistance in peripheral tissues (Xin *et al*. 2016).Of the 89 bulk samples, the HbA1c level information is available for 77 individuals, which includes 51 healthy and 26 diabetic individuals. Xin *et al*. (2016)used single-cell RNA-seq to measure the transcriptomes in 1492 human islet cells from 12 healthy and 6 T2D organ donors. They focused on four major endocrine cell types (alpha, beta, delta and gamma). The scRNA-seq data for healthy samples was composed of 354 alpha, 194 beta, 28 delta and 33 gamma cells while the scRNA-seq data for T2D samples was composed of 532 alpha, 278 beta, 21 delta and 52 gamma cells.

We used the scRNA-seq datasets to capture cell-type proportions in the bulk samples. Figure 3 represents estimated beta cell-type proportions versus HbA1c levels (after outlier removal) as well as p-values for a negative Spearman correlation test between the beta cell-type proportions and HbA1c levels. Based on the p-values, DWLS, CSCD, SVR and CIBERSORTx were successfully able to detect a strong negative relationship between beta proportions and HbA1c levels. It is remarkable that SVR and OLS reported lower estimated values than the other methods.

**Figure 3:**
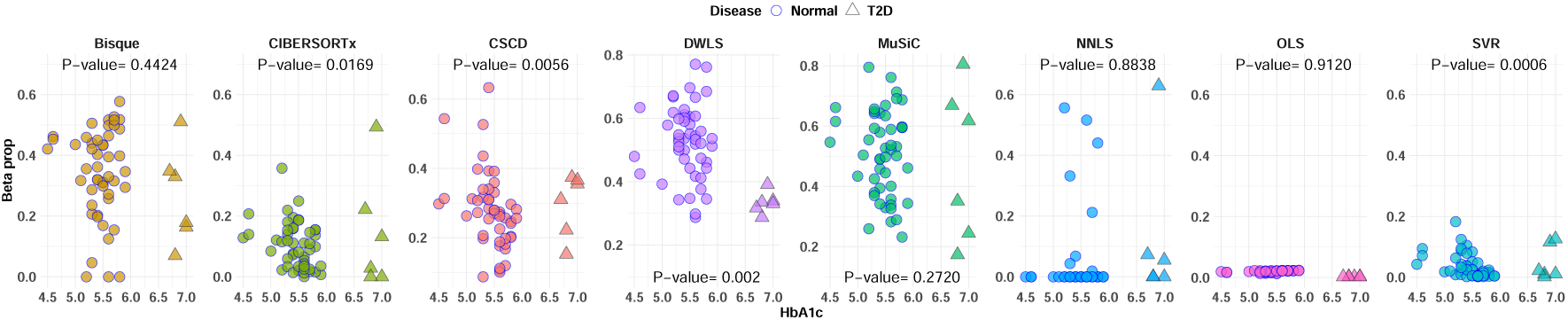
HbA1c levels versus estimated beta proportions in bulk samples in the pancreatic islets data (Fadista *et al*. 2014). The p-values were obtained using a negative Spearman correlation test between beta proportions and HbA1c levels.

### 3.4 Evaluation on cortex tissue data

As another application, we benchmarked all these decomposition methods using expression data collected from dorsolateral prefrontal cortex (DLPFC). This dataset was generated by the Rush Alzheimer’s Disease (AD) Center (Mostafavi *et al*. 2018) and includes 636 postmortem bulk RNA-seq samples. The scRNA-seq data were collected from 24 of the 636 individuals. The scRNA-seq data include 162767 cells that have already been clustered into eight clusters, as provided in Table 1. To manage the memory limit available on our system, we randomly selected eight individuals and this reduced the number of cells to 60388 cells, see Table 1. We remark that most scRNA-seq data consist of only a few samples. Similarly, Bisque (Jew *et al*. 2020) developers also used eight individuals to study the performance of their method. We further, similar to Bisque, sampled 25% of the nuclei in the scRNA-seq data to accommodate CIBERSORTx’s uploading limitation, which currently is 1GB.

Figure 4 represents a jitter plot of the estimated cell-type proportions using the different methods for the eight individuals common in both bulk and scRNA-seq samples. Each panel represents real and estimated values for the labelled cell type. From Figure 4, it is obvious that all methods except Bisque and CSCD either under-estimated or over-estimated the proportions. CIBERSORTx, MuSiC and NNLS under-estimated Peri cell-type proportions while SVR, DWLS and OLS over-estimated them. All the methods except Bisque and CSCD under-estimated Exc proportions. Quantitative comparison of data is summarized in Table 4 and shows the corresponding R, MSE and MAE values. From this table, we observe that all methods led to poor correlation results. CIBER-SORTx led to a negative correlation, and CSCD, DWLS and SVR led to a bit higher correlation. Looking at MSE and MAE measurements in Table 4, it is observed that CSCD outperformed all the other tools. Bisque and DWLS were the second and third top methods, respectively.

**Table 4:** Performance measures of different cell-type estimation methods based on eight bulk samples in the cortex data.

| Measure | CSCD | Bisque | CIBERSORTx | MuSiC | SVR | NNLS | DWLS | OLS |
| --- | --- | --- | --- | --- | --- | --- | --- | --- |
| R | 0.1250 | 0.0179 | -0.0598 | 0.0003 | 0.1499 | 0.0206 | 0.1577 | 0.0486 |
| MSE | 0.0014 | 0.0015 | 0.0749 | 0.0663 | 0.0231 | 0.1230 | 0.0129 | 0.0274 |
| MAE | 0.0234 | 0.0266 | 0.1818 | 0.1812 | 0.1235 | 0.2188 | 0.0881 | 0.1340 |

**Figure 4:**
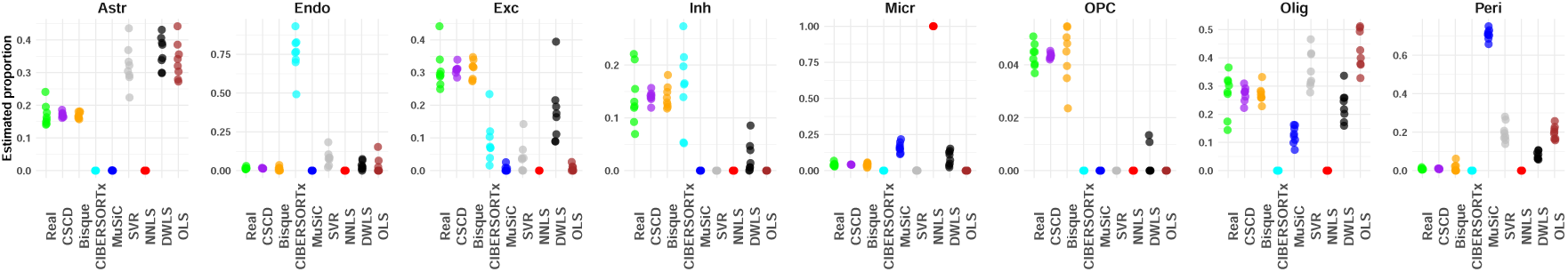
Jitter plot of estimated cell-type proportions for eight bulk samples in the cortex tissue data. Each panel represents estimated values for the labelled cell type. Different colors represent different estimation methods.

We further applied these decomposition methods to estimate cell-type proportions in the remaining 628 individuals based on expression levels of the eight distinct individuals in the scRNA-seq data. Figure 5 represents the observed scRNA-seq proportions as well as estimated proportions using the eight methods. The Figure illustrates that both CSCD and Bisque were successfully able to preserve the distribution of the scRNA-seq data, and produced estimates in bulk data that were significantly close to the scRNA-seq observations.

**Figure 5:**
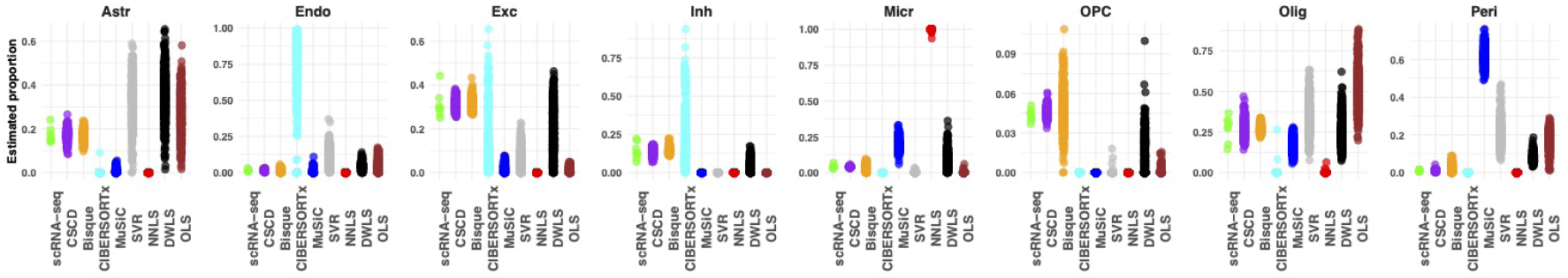
Jitter plot of estimated cell-type proportions for 628 bulk samples in the cortex tissue data. Each panel represents estimated values for the labelled cell type. Different colors represent different estimation methods.

To determine cell-specific decomposition accuracy, we measured association between neuron proportions (the sum of Exc and Inh proportions) and each individual physician cognitive diagnostic category at time of death (cogdx). Neural death is a hallmark symptom of AD (Yankner 1996), and a negative association between cogdx and neuron proportion is expected (Jew *et al*. 2020). The cogdx provides a semiquantitative measure of dementia severity ranging from 1 (no cognitive impairment) to 5 (a confident diagnosis of AD by physicians). Figure 6 represents estimated neuron proportion according to different cogdx levels. Based on this figure alone, it is difficult to make any inference on association between cogdx and proportions. We further validated the negative relationship between codex levels and neuron proportions. Because the cogdx variable has ordered discrete levels from 1 to 5, we used the Jonckheere-Terpstra test (Jonckheere 1954). We included the resulting p-values in Figure 6. Based on the p-values and the significance level 0.01, we observe that all methods except NNLS were able to identify the expected negative association between neuron proportions and cogdx levels. It is notable that SVR, OLS, NNLS and MuSiC reported lower estimated proportion values than the other methods.

**Figure 6:**
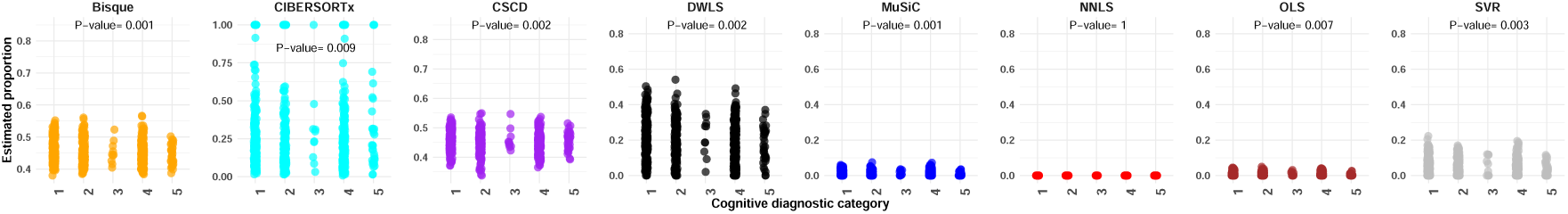
Jitter plot of estimated neuron proportions for 628 bulk samples in the cortex tissue data, according to individuals physician cognitive diagnostic category at time of death. Each panel represents neuron proportions estimated by one of the eight different methods.

## 4 Discussion and Concluding Remarks

In this paper, we presented a reference-based decomposition method. Our primary goal was to improve on Bisque. We presented the CSCD approach and illustrated the method performance by analyzing several datasets. CSCD outperformed most of the existing methods evaluated in this study. By using both bulk and scRNA-seq datasets that were collected from the same individuals, we observed that CSCD is able to capture the expected cell-type proportions.

The method is executable by the inclusion of all genes in the bulk and/or scRNA-seq data in the analysis. However, it may lead to imprecise results, as some genes may appear in most (or all) of cell-types categories. To overcome this issue, one may select genes to be included in the analysis based on a preferred marker gene list. This is indeed the way scRNA-seq data are often prepared. Clusters in scRNA-seq data are constructed based on a pre-collected list of identified marker genes. Genes not on the original list are then assigned to clusters using some probabilistic approaches such as the nearest neighbor or t-SNE clustering, which could in turn be a source of noise. If such an original marker gene list was available in addition to a given scRNA-seq dataset, one could just base the analysis on the makers on the given list. Otherwise, we suggest identifying marker genes using one of the existing approaches in the literature. We used the MAST approach (Finak *et al*. 2015) implemented in the `FindAllMarkers` function of the `Seurat` package (Hao *et al*. 2021), and our results indicated that this approach works well.

CSCD is similar to Bisque in the sense that it is a transformation-based approach and standardizes gene expression levels but its strength is the ability to successfully incorporate the covariance structures in both scRNA-seq and bulk RNA-seq datasets into the analysis. This leads to a more accurate transformation and more accurate cell-type estimates. Also, CSCD applies the linear shrinkage estimation of the covariance matrices, accounting for the collinearity issue, which is common in every multivariate large-scale data analysis.

CSCD and Bisque were robust to changes in data sources. For example, looking at the cortex tissue data, CSCD was the top method, and Bisque and DWLS were the second and third top methods, respectively. Looking at the pancreatic islets data, CSCD and Bisque were the first and second top methods, with SVR being the third. Both Bisque and CSCD rely on transformations to match the distribution of both single-cell expression and bulk expression data. This requires enough samples in both datasets. Having a small number of individuals may lead to unreliable results. Also, both Bisque and CSCD preserve the distribution of data in scRNA-seq. Both methods assume that the resulting single-cell-based estimates of cell-type proportions accurately reflect the true cell-type proportions of interest. If cell-type proportions captured by single-cell experiments differ significantly from the true physiological distributions, the accuracy of Bisque and CSCD may decrease. Users are advised to be cautious in case of existence of a known significant bias in the single-cell measurements of a tissue of interest. For more discussion, readers may refer to Jew *et al*. (2020).

## Acknowledgements

The author would like to thank the Bioinformatics Division of the Bureau of Food Surveillance and Science Integration of Health Canada for the computing facilities and support. The author also wishes to thank Brandon Jew for providing useful information about Bisque as well as the DLPFC data. The embryonic stem cells data (both bulk and scRNA-seq) were downloaded from the Gene Expression Omnibus (GEO) database with accession number GSE75748. The pancreatic islets data of Segerstolpe *et al*. (2016) were downloaded from the ArrayExpress (EBI) database with accession numbers E-MTAB-5061 (bulk) and E-MTAB-5060 (scRNA-seq). Processed bulk data of Fadista *et al*. (2014) and scRNA-seq data of Xin *et al*. (2016) were download from https://xuranw.github.io/MuSiC/data. The DLPFC data are available on Synapse (10.7303/syn3219045). ScRNA-seq data (syn23554292), bulk RNA-seq data (syn3505720), and phenotypes (syn3191087) are available under controlled use conditions set by human privacy regulations. A data use agreement is required to access these data. This research was enabled in part by support provided from Health Canada, University of Ottawa and Compute Canada (www.computecanada.ca).

